# Aspartic proteases are abundant and active in acidified wound fluid

**DOI:** 10.1101/2024.06.30.601389

**Authors:** Elany Barbosa da Silva, Meredith J. Crane, Lawrence Liu, Danielle J. Gelsinger, Alexander R.D. Jordon, Robin L. McKinney, Craig P. Eberson, Amanda M. Jamieson, Anthony J. O’Donoghue

**Author notes:** Corresponding authors &.

## Abstract

Wound healing necessitates a balance between synthesis and breakdown of extracellular matrix components, which is tightly regulated by proteases and their inhibitors. Studies have shown that treatment of poorly healing wounds with acid results in improved healing. In this study, we systematically evaluated changes in proteolytic activity of murine wound fluid upon acidification. A library of 228 synthetic peptides served as reporters of protease activity at pH 7.4, pH 5.0 and pH 3.5. The peptide digestion patterns differed at each pH, revealing that proteases active at pH 7.4 are inactivated at pH 3.5. Notably, aspartic acid proteases emerged as the dominant active enzymes at pH 3.5 and their activity was inhibited by pepstatin. Using a fluorogenic substrate, we quantified aspartic protease activity across varying pH levels and demonstrated optimal activity between pH 3.0 and 3.8. This activity was detectable as early as one day post-injury and persisted over the following ten days. Importantly, human wound fluid exhibited the same activity profile, validating the mouse model as a relevant system for studying acid-mediated wound healing processes.

## Introduction

The successful healing of adult skin involves carefully coordinated phases, including hemostasis, inflammation, proliferation, and remodeling. Disruption of any of these stages can derail the process leading to chronic non-healing wounds (1, 2). Elderly individuals and those with diabetes are at especially high risk of developing poorly healing wounds, requiring an estimated $28B annually in healthcare associated expenditures (3, 4). With over 20% of the US population expected to be age 65 or older by the year 2030, and the number of people being diagnosed with type 2 diabetes increasing year-after-year, the medical and financial costs associated with poorly healing wounds are projected to increase accordingly. Therefore, it is important to better understand the numerous molecular players that coordinate the stages of tissue repair at steady state and in chronic settings to inform new avenues of treatment for poorly-healing wounds. Proteases play crucial roles in all the stages of the wound healing process, from the initial hemostasis stage through extracellular matrix remodeling. To date, most of the well-studied proteolytic activities in wound healing are metalloproteinases and serine proteases. Matrix metalloproteinases-2 (MMP-2) and -9 (MMP-9) degrade the components of the extracellular matrix, assist in regenerating injured tissues, and help cellular migration to the wound area (5). Serine proteases such as cathepsin G, elastase and urokinase-type plasminogen activator (uPA) are expressed by immune cells and endothelial cells in the wound microenvironment, and they are mainly involved in the re-epithelialization process and degradation of growth factors. Of note, each of these enzymes are optimally active at neutral pH (1).

The typical pH range of human skin is between 4.2 and 5.6, which creates an acidic environment that serves as a protective barrier against bacterial overgrowth (6). In contrast, acute wounds exhibit a neutral to alkaline pH spanning from 6.5 to 8.5, and chronic wounds tend to fall within the range of 7.2 to 8.9 (6). Chronic wounds have been associated with increased protease levels that are influenced by the pH milieu of the wound (7).

Reagents that can lower wound pH, such as citric acid (CA) and acetic acid have been used to promote wound healing. These treatments have been shown to positively correlate with accelerated wound healing (8–11). In a notable case study of a patient with multiple leg ulcers, 33 applications of a 3% CA, resulted in complete healing (12). This outcome stands in contrast to the ineffectiveness of conventional therapies whereby amputation had been recommended to this patient prior to CA treatment. In another case study, daily application of 3% CA for 11 days achieved resolution of a nonhealing tuberculous sinus. Notably, the sinus had shown resistance to all conventional antimicrobial therapies and local care interventions (8). In a larger study consisting of 115 patients with diabetic foot ulcers infected with diverse bacteria, treatment using a CA gel resulted in 106 patients experiencing healing (13). An acidic environment is generally unfavorable to bacteria, so the treatments are thought to reduce infection. Studies have shown that CA treatment results in a higher percentage of re-epithelialization, a slenderer epithelial layer, improved wound contracture, and elevated levels of collagen deposition in non-infected wounds (14). However, the mechanism of how acidification of wounds affects proteolysis during healing, has not been fully elucidated.

Enzymes that are well characterized in the wound healing setting, like MMP-2, MMP-9, and uPA are unlikely to be active in acidic conditions. Therefore, we sought to assess protease activity in an acidified wound. We utilized a peptide digestion assay known as Multiplex Substrate Profiling by Mass Spectrometry (MSP-MS) to determine the peptide cleavage profile generated by wound fluid proteases at three pH conditions (as illustrated in **Figure 1**). More specifically, this assay consists of a library of 228 strategically designed 14-mer peptides. Upon addition of a biofluid sample, cleavage of any of the 2,964 peptide bonds (228 peptides x 13 bonds) in the library by proteases can be detected using tandem mass spectrometry. This method has previously been used to discover new protease activities in biological samples such as cancer cell secretions (15), plasma (16), gastric juice (17), and pancreatic cyst fluid (18). We anticipated that peptides in this library would be cleaved by proteases in the wound fluid when assayed at neutral pH as these proteases have already been well characterized. We hypothesized that the protease activity in the same wound fluid samples would be reduced when assayed at acidic conditions, since the optimum pH of MMPs and serine proteases is close to neutral pH (19–21). However, we were surprised to find that acidification of wound fluid activates aspartic acid proteases and the total number of peptides that are cleaved is higher at pH 3.5 when compared to pH 7.4. In this study, we performed an in-depth characterization of the enzyme activity derived from aspartic acid proteases in murine wound fluid and then revealed that similar activity exists in human wound fluid. This study sheds light on the changes in protease activity upon acidification of wounds.

**Figure 1:**
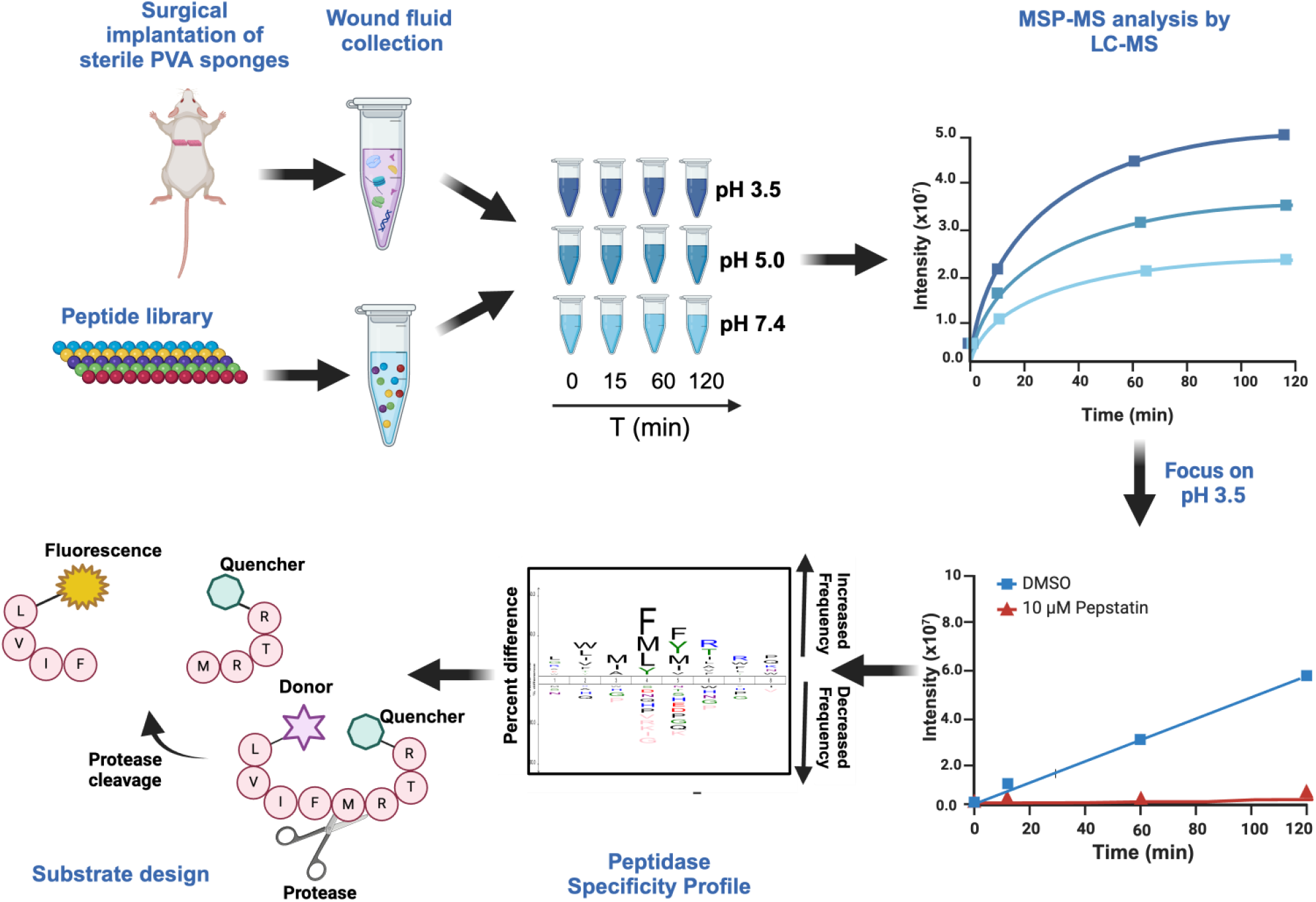
Overview of the substrate profiling studies with murine wound fluid at pH 3.5, 5.0, and 7.4 and the development of fluorogenic reporter substrates.

## Methods

### Mouse models and wound fluid retrieval

All animal studies were approved by the Brown University Institutional Animal Care and Use Committee (protocol #22-04-0001) and performed in accordance with the Guide for the Care and Use of Animals of the National Institutes of Health. C57BL/6J mice were bred in-house. Male mice aged 8-12 weeks were used for studies. Wound fluid samples were collected using the subcutaneous polyvinyl alcohol (PVA) sponge implantation model, which allows for the retrieval of wound fluid with minimal manipulation (22–24). The model recapitulates the phases of acute wound healing, including inflammatory, angiogenic, and fibrotic responses seen in soft tissue wounds, for up to 2 weeks following implantation. For PVA sponge implantation surgery, mice were placed under anesthesia and analgesia using ketamine (80-100 mg/kg) and xylazine (8-10 mg/kg), respectively. The dorsum was shaved and cleaned with povidone-iodine solution and 70% ethanol. Under sterile conditions, a 2 cm full-thickness incision was made along the dorsal midline. Six sterile 1 cm x 1 cm x 0.5 cm PVA sponges were implanted into subcutaneous pockets on either side of the midline. The incision was then closed using stainless steel surgical clips. In this model, fluid that accumulates in response to the injury is captured by the sponges, which were then removed from the mice at various time points (day 1, 3, 5, 7, and 10). To isolate wound fluid, PVA sponges were placed into the barrel of a 5 mL syringe that was nested in a 15 mL culture tube. The tube was centrifuged at 500 x g for 10 minutes, allowing fluid to be drawn out of the sponges into the collection tube. Cell-free fluid was stored at -80°C (25).

### Spinal fusion surgical wound drainage sample collection

All human studies were approved by the Lifespan Institutional Review Board (IRB #_**1135267**) and informed consent was obtained from the patient or guardian. Drainage samples were collected from idiopathic (AIS) or neuromuscular (NM) scoliosis patients recovering from spinal fusion surgery. As part of the standard protocol for pediatric spinal fusions, drains were placed either subcutaneously for AIS patients or in the submuscular space for patients with NM scoliosis, the latter group usually undergoing a multi-layered closure with muscle flaps due to well-documented healing issues in this group. While drains for the AIS patients were usually removed by the second or third postoperative day, the drains for the NM patients usually had higher output due to their location below the muscle layer and thus remained in place for the entire hospitalization in most cases, resulting in longer sampling time. Daily samples were collected from the effluent of wound drains placed at the surgical incision. Once collected, samples were kept on ice until the time of processing. Drainage fluid was centrifuged at 300 x g, and the cell-free supernatant was stored at -80°C.

### BCA assays

Protein in mouse and human wound fluid samples were quantified with the BCA Protein Assay kit (Thermo Fisher Pierce) using the 96-well plate protocol recommended by the manufacturer. A serial dilution of bovine serum albumin standards was evaluated in parallel with a 1 in 10 and 1 in 50 dilution of each wound fluid sample. 25 µL of the standard or unknown was then incubated with 200 µL of the working reagent, which was included in the assay kit, at 37°C for 30 minutes. The plate was then read at 562 nm using a Synergy HTX (Biotek) plate reader. A standard curve was created, and the dilution of each sample that fell within this range was computed and used to determine the total protein concentration in the sample. These stock concentrations were then used to normalize the activity to protein concentration.

### Protease cleavage profile by MSP-MS

Multiplex substrate profiling by mass spectrometry (MSP-MS) utilizes a library of 228 tetradecapeptides designed to contain all neighbor and near-neighbor cleavage sites (26). Wound fluid was extracted from 3 mice on day 5 post injury and was diluted 110-fold in each of the following assay buffers: Assay Buffer 1: 20 mM sodium citrate pH 3.5 and 100 mM sodium chloride (NaCl); Assay Buffer 2: 20 mM sodium acetate pH 5.0 and 100 mM NaCl; and Assay Buffer 3: Dulbeccòs phosphate buffer saline (D-PBS) pH 7.4 (Gibco 14040-133). In parallel, the library of peptides was diluted 21.9-fold in the same three buffers such that the concentration of each peptide was 1 μM. Then an equal volume of wound fluid in assay buffer was mixed with peptides in the same assay buffer such that the final concentration of each peptide is 0.5 μM.

Assays were incubated at 25°C for 15, 60, and 120 minutes. At each time point, 20 µL of the reaction mixture was removed and quenched by the addition of urea (Sigma product #SLBT1918) to a final concentration of 4 M. Samples were immediately stored at -80°C. Control reactions consisted of mice wound fluid samples pre-incubated with 4 M urea to inactivate the enzymes prior to the addition of the peptide library. All samples were desalted using a C18 column and dried in a vacuum centrifuge as outlined previously (27). All assays were performed in quadruplicate reactions.

In a follow-up study, wound fluid from day 5 and day 10 was diluted 110-fold in Assay Buffer 1 containing 20 μM of pepstatin or 0.2% dimethyl sulfoxide (DMSO) as the vehicle control. The reaction was incubated for 60 minutes and then was mixed with an equal volume of the peptide library diluted in Assay Buffer 1. Assays were incubated at 25°C for 15, 60, and 120 minutes and 20 µL of the reaction mixture was removed and quenched by the addition of urea (Sigma product #SLBT1918). The control reaction (0 minutes) consisted of wound fluid pre-incubated with 4 M urea prior to addition of the peptide library.

For each MSP-MS assay sample, ∼0.4 µg of peptides was injected into a Q-Exactive Mass Spectrometer (Thermo) equipped with an Ultimate 3000 HPLC. Peptides were separated by reverse phase chromatography on a C18 column (1.7 um bead size, 75 um x 25 cm, 65°C) at a flow rate of 300 nL/min using a 60-min linear gradient from 5 to 30% B, with solvent A: 0.1% formic acid in water and solvent B: 0.1% formic acid in acetonitrile. Survey scans were recorded over a 150–2000 m/z range (70,000 resolutions at 200 m/z, AGC target 3×10^6^, 100 ms maximum). MS/MS was performed in a data-dependent acquisition mode with HCD fragmentation (28 normalized collision energy) on the 12 most intense precursor ions (17,500 resolutions at 200 m/z, AGC target 1×10^5^, 50 ms maximum, dynamic exclusion 20 s). Data was processed using PEAKS 8.5 (Bioinformatics Solutions Inc.). MS^2^ data were searched against the tetradecapeptide library sequences with decoy sequences in reverse order. A precursor tolerance of 20 ppm and 0.01 Da for MS^2^ fragments was defined. No protease digestion was specified. Data were filtered to a 1% peptide level false discovery rate with the target-decoy strategy. Peptides were quantified with label free quantification, and data were normalized by median and filtered by 0.3 peptide quality. Missing and zero values were imputed with random normally distributed numbers in the range of the average of the smallest 5% of the data ± SD.

### Aspartyl proteases activity and inhibition assays

Activity of murine wound fluid samples from day 1, 3, 5, 7, and 10 was measured by monitoring the cleavage of the fluorescent substrate, Mca-Gly-Lys-Pro-Ile-Leu-Phe-Phe-Arg-Leu-Lys(DNP)-dArg-NH2 (shortened to GKPILFFRL) (CPC Scientific) using a Synergy HTX (Biotek) fluorimeter. A 2 mM stock solution of this substrate was prepared in DMSO. All assays were performed in black flat-bottom 384-well plates (Greiner, catalog #784900), in 30 μL of Assay Buffer 1 supplemented with 2 mM dithiothreitol (DTT), 0.01% Tween-20, 20 µM GKPILFFRLK, and 1 in 110 dilution of wound fluid. Prior to addition of the substrate, enzyme was incubated with 20 μM pepstatin or 0.2% DMSO for 1 hour. Following the substrate addition, the change in fluorescence was recorded for 2 hours. Enzymatic activity in relative fluorescence per second (RFU/sec) was calculated from the initial rates of the reaction. All assays were run in triplicate wells. Using the total protein content in the assay, the results were then normalized to RFU/s per µg of protein (RFU/s/µg), to allow for comparison across samples. For human wound fluid assays, individual samples from postoperative day 1, 2, and 3 were evaluated using the fluorescent substrate GKPILFFRL in the presence and absence of 20 μM pepstatin and employing the same protocol described previously for mouse samples.

### Protease activity assays across a pH range

Pooled mouse wound fluid samples from day 5 and 10 were assayed with 20 µM GKPILFFRL in sixteen different citrate phosphate buffers ranging from pH 2.2 to pH 8.0, and two Tris buffers at pH 8.6 and 9.5. All buffers consisted of 100 mM citrate phosphate or 100 mM Tris-HCl. Experiments were performed in triplicate wells using a Synergy HTX (Biotek) fluorimeter.

### Proteases activity in human wound fluid and inhibition assays

Activity of human surgical wound fluid samples from one AIS and three NM patients was measured by monitoring the cleavage of the fluorescent substrates, GKPILFFRL, Mca-Arg-Pro-Lys-Pro-Val-Glu-Nva-Trp-Arg-Lys(DNP)-DArg-NH2 (shortened to RPKPVEvWR) (CPC Scientific), and MCA-Pro-Leu-Gly-Leu-Dap(DNP)-Ala-Arg-NH2 (shortened to PLGLdAR) (CPC Scientific), using a Synergy HTX (Biotek) fluorimeter. A 2 mM stock solution of the substrates was prepared in DMSO. All assays were performed in black flat-bottom 384-well plates (Greiner, catalog #784900), in 30 μL using three different pH buffers: 1) 20 mM sodium citrate, 100 mM NaCl, pH 3.5, supplemented with 2 mM dithiothreitol (DTT), 0.01% Tween; 2) 20 mM sodium acetate pH 5.0, 100 mM NaCl, 2 mM DTT, 0.01% Tween; and 3) Dulbeccòs phosphate buffer saline pH 7.4 (Gibco 14040-133). All substrates were tested at 20 µM, using 1 in 110 dilution of wound fluid in each assay buffer. For inhibitor studies, enzyme was pre-incubated for 1 hour with 20 μM pepstatin, 1 mM Ethylenediaminetetraacetic Acid (EDTA), 100 μM marimastat, 10 μM *trans*-Epoxysuccinyl-L-leucylamido(4-guanidino)butane (E64), 1 mM 4-(2-aminoethyl)benzenesulfonyl fluoride hydrochloride (AEBSF), and 10 μM carfilzomib. Following the substrate addition, the change in fluorescence was recorded for 2 hours. Enzymatic activity in relative fluorescence per second (RFU/sec) was calculated from the initial rates of the reaction. All assays were run in triplicate technical replicates and DMSO was used as the vehicle control. Using the total protein content in the assay, the results were then normalized to RFU/s per µg of protein (RFU/s/µg), to allow for comparison across samples.

## Results

### Global Protease Substrate Profiling revealed pH-specific activities

To uncover the protease activities in murine wound fluid at different pH conditions, wound fluid was pooled from three mice 5 days post PVA implantation and incubated with an equimolar mixture of 228 14-mer peptides in buffers that varied by pH. For the normal conditions, pH 7.4 was chosen as this corresponds to the pH of the murine wound fluid. Assays were also performed in pH 5.5 and pH 3.5 buffers to mimic a mildly acidic and moderately acidic environment, respectively. Following two hours incubation with the peptide mixture, cleaved products were found in all pH conditions, however, the site of peptide cleavage often differed. Only one peptide in the library, consisting of the sequence, YTRLNGEAVnFnSK, was cleaved by proteases at all pH conditions, however the location of the cleavage site differed at different pHs. At pH 7.4, proteases cleaved between nS and SK. However, at pH 5.0 cleavage occurred between Fn and SK, while at pH 3.5 cleavage occurred between nF and Fn (**Figure 2**). As there is no overlap between the cleavage sites generated at pH 7.4 and pH 3.5, we hypothesize that proteases active at one pH are inactive at the other. In addition, protease activity at pH 5.0 is likely due to some of the neutral and acid proteases retaining activity at this intermediate condition.

**Figure 2:**
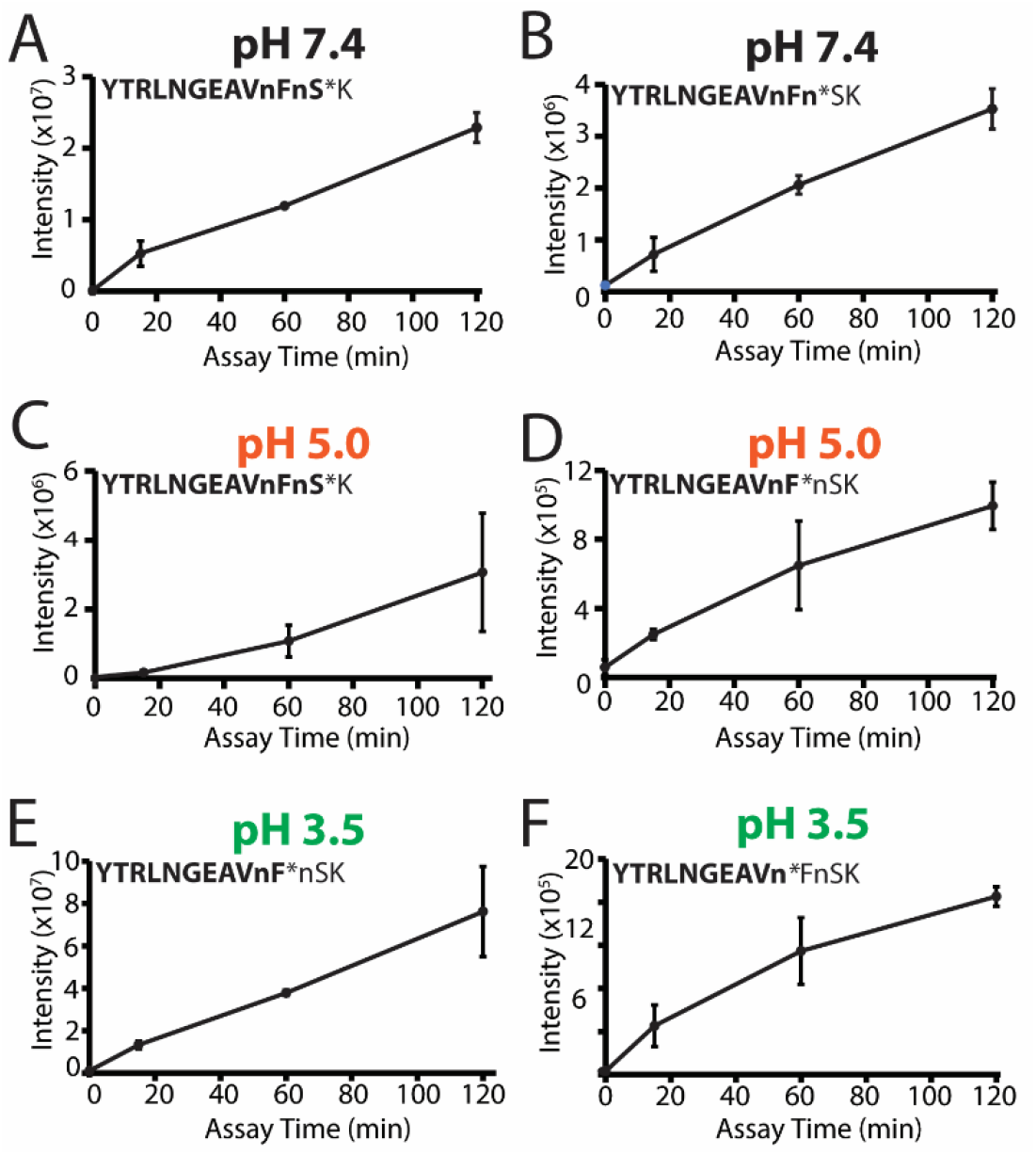
pH dependent cleavage of YTRLNGEAVnFnSK by murine wound fluid proteases over time. **A**. Peptide was cleaved between S and K at pH 7.4 and the cleavage site is indicated by *. The cleavage product (highlighted in bold font) was quantified by LC-MS/MS after 0, 15, 60 and 120 minutes incubation. **B.** A second cleavage product detected and quantified at pH 7.4. **C**-**D**. Cleavage products quantified at pH 5.0. **E-F** Cleavage products quantified at pH 3.5. All assays were performed in four technical replicates using a pooled wound fluid sample from three mice. Lowercase n corresponds to norleucine.

When wound fluid was assayed at pH 7.4, proteases cleaved at 29 sites. The distribution of these cleavage sites across the 14-mer peptides, as illustrated in **Figure 3A**, revealed that 16 of the 29 cleavage sites occurred between the 12^th^ and 13^th^ residues. The proteases responsible for cleavage at this site are categorized as di-carboxypeptidases, as these enzymes remove two amino acids from the carboxy terminus. In addition, four peptides were cleaved between the 13^th^ and 14^th^ amino acids indicating the presence of mono-carboxypeptidases while the remaining cleavage occurred near the N-terminus or in the middle of the peptides.

**Figure 3:**
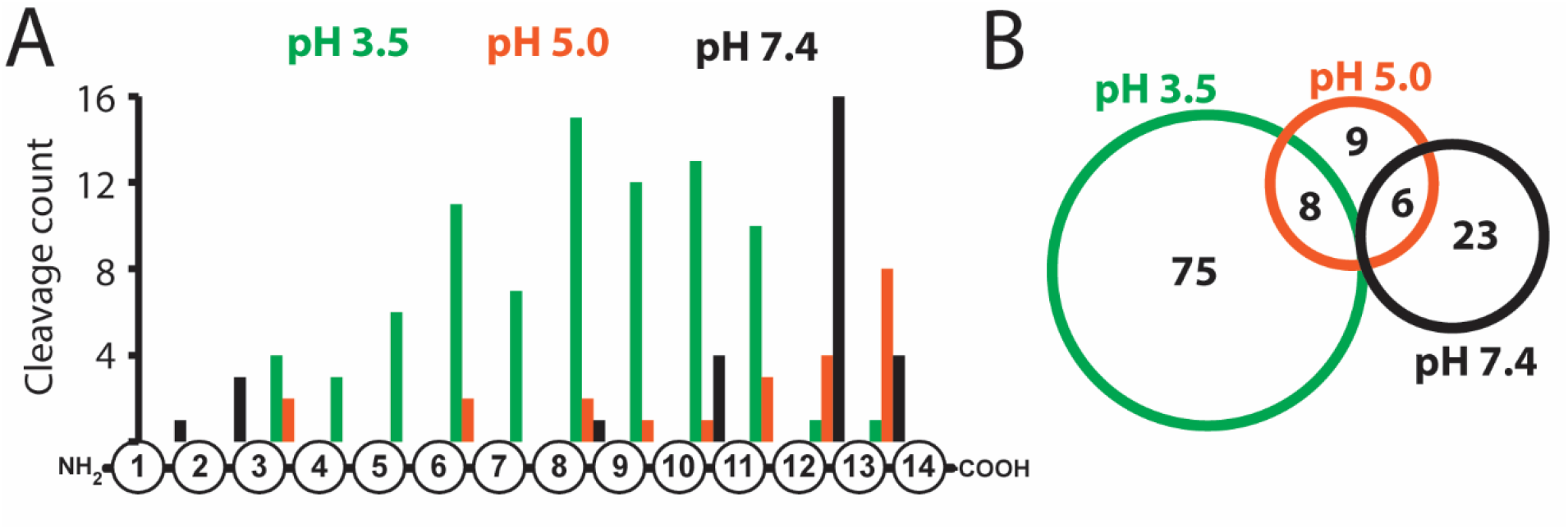
Quantitative multiplex substrate profiling of murine wound fluid. **A**. Distribution of murine wound fluid protease cleavage sites within 14-mer peptides. **B**. The Venn diagram shows the number of cleavage sites shared between the three different pH conditions.

At pH 5.0, the distribution of cleavage sites differs to that seen at pH 7.4 conditions. Here, 8 out of the 23 cleavage sites were between the 13^th^ and 14^th^ residues while only 4 peptides were cleaved between the 12^th^ and 13^th^ amino acids. These data reveal that the mono-carboxypeptidase activity is stronger than the di-carboxypeptidase activity at pH 5.0. Finally, at pH 3.5, we were surprised to detect 83 cleaved peptides, generated by proteases in the wound fluid. The distribution of these sites was spread been the 3^rd^ and 12^th^ amino acids and only 1 peptide was cleaved between the 12^th^ and 13^th^ residues and another one between the 13^th^ and 14^th^ residues. This distribution indicates that most proteases that are active at pH 3.5 can be defined as endoproteases, as these enzymes preferentially cleave away from the N- and C-termini.

We next assigned each of the potential 2,964 cleavage sites (228 peptides x 13 peptide bonds) within the library a unique identifier and then determined which of them were cleaved by wound fluid proteases across the three pH conditions. We found that there was no overlap between the sites cleaved by proteases at pH 3.5 and pH 7.4, while there were 8 sites in common between pH

3.5 and pH 5.0, and 6 sites in common between pH 5.0 and pH 7.4 (**Figure 3B**). In addition, there are proteases that are active at pH 5.0 and not at the more acidic and neutral conditions since there were 9 cleavage sites that were generated only at pH 5.0. These data reveal that acidification of wound fluid results in the activation of distinct acid-acting proteases and inactivation of the neutral-acting proteases.

We next generated a substrate specificity profile at each pH condition to illustrate the similarities and differences in the murine protease activity with changing pH. At pH 3.5, peptides were cleaved frequently when Phe, Leu, Nle (norleucine, n) and Tyr were in the P1 position while Phe, Nle, Tyr and Ile were favored at P1′. At other positions Arg at the P2′ and P3′ was frequently observed while non-polar amino acids were found with highest frequency at P2 and P3 (**Figure 4A**). At pH 5.0, non-polar amino acids were generally found at P3, P1 and P1′, while Pro was favored at P4′ and Lys and Arg were favored at P1′ and P2, respectively (**Figure 4B**). At pH 7.4 positively charged amino acids such as Arg and Lys were favored at P1′ while non-polar amino acids in addition to Arg were also frequently found at P1 (**Figure 4C**). These findings underscore the diverse protease activities present in wound fluids across a broad pH range.

**Figure 4:**
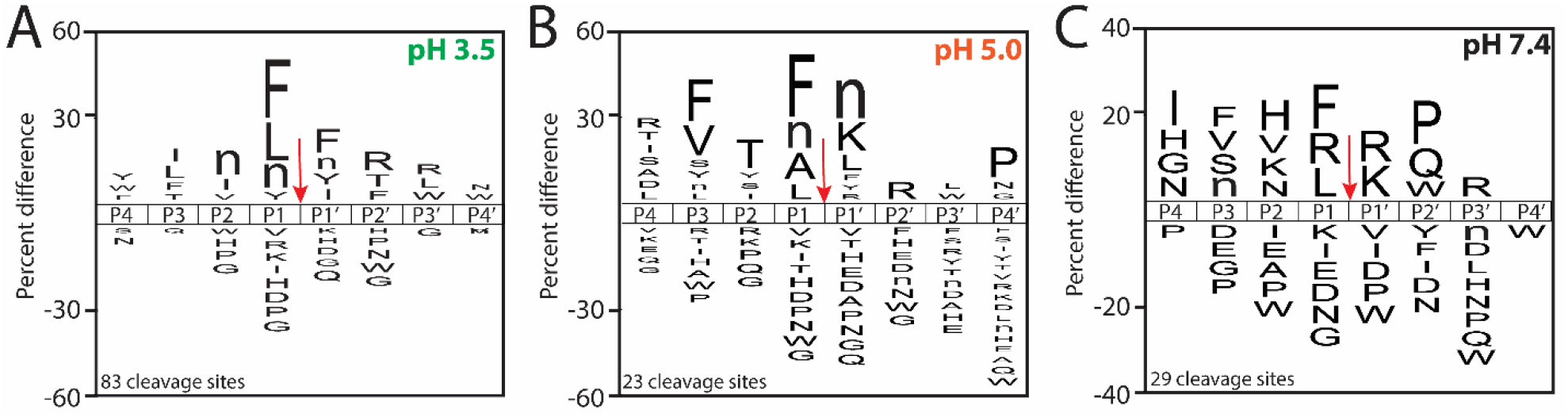
Icelogo plots illustrating the cleavage profiles. Wound fluid samples taken 5 days after injury were assayed at pH 3.5, 5.0 and 7.4. Data represents a pooled wound sample from 3 mice assayed in quadruplicate reactions. The red arrow indices the site of cleavage.

Following these studies at different pH conditions, we then focused our attention on the protease activities present at pH 3.5 since these enzymes are likely to be active in acid-treated wounds. We first sought to determine if differences in proteases activity exists in murine wound fluid taken 5 days and 10 days after injury. Wound fluid from Day 10 was subjected to MSP-MS analysis in pH 3.5 assay buffer. In total, 108 cleavage sites were quantified, of which 81 were identical to those found in the Day 5 sample (**Figure 5A**). Most of the additional 27 cleavage sites detected on Day 10 were also observed in the Day 5 sample, however the cleavage products were not sufficiently abundant to pass our stringent statistical testing. These data reveal that the same acid-acting proteases exist in wound fluid on Day 5 and Day 10, but the activity is higher on Day 10. To evaluate further, the cleavage products generated in each dataset were directly compared. One of the most efficiently cleaved substrates was PHWQRVIFFRLNTP where cleavage occurs between FF. We quantified the time-dependent accumulation of both the N-terminal fragment, PHWQRVIF (**Figure 5B**) and the C-terminal fragment, FRLNTP (**Figure 5C**) by proteases present in Day 5 and 10 wound fluid. In both assays, the products were generated more rapidly by proteases in the Day 10 sample.

**Figure 5:**
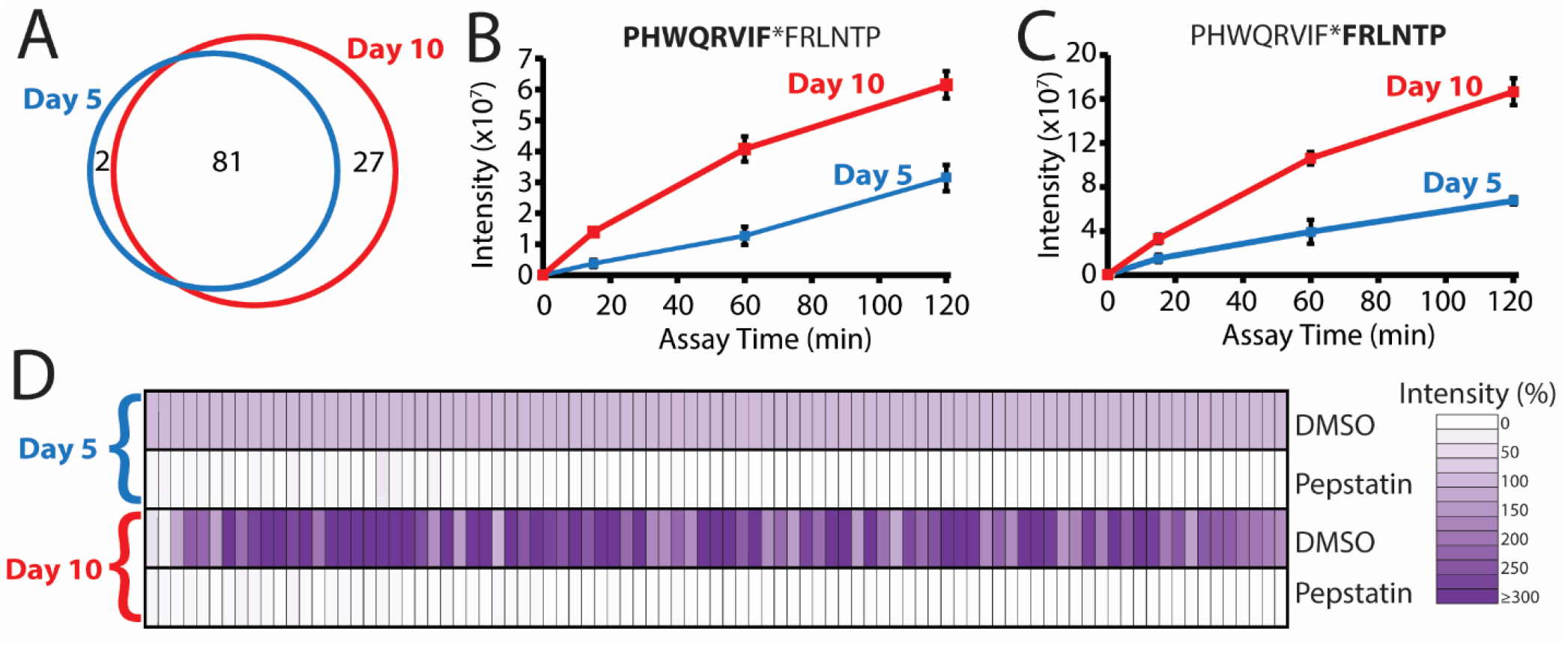
Comparation of proteolytic activity in wound fluid by MSP-MS 5- and 10-days post- injury. **A.** Number of cleavage sites detected at pH 3.5 in wound fluid 5- and 10-days post-injury. **B-C.** Quantification of the time-dependent accumulation of select N-terminal and C-terminal fragments. **D**. Normalized intensity of proteolytic cleavage products in the presence and absence of pepstatin. Each rectangle represents the average intensity of a cleaved peptide product (n = 4 technical replicates) normalized to the intensity of the same cleaved peptide that was quantified in the Day 5 sample. Lighter color indicated that the cleaved peptide decreased in intensity while darker color indicated that the peptide increased.

To compare all cleavage sites, we first normalized the intensities of each cleavage product found after 120 minutes incubation in the Day 5 wound fluid sample and then quantified the relative abundance of the same cleavage product generated by proteases in the Day 10 sample. We show that almost all peptide products are higher in intensity following incubation with Day 10 wound fluid, with some being more than 3-times higher (**Figure 5D**).

In this assay, we also pre-incubated the wound fluid with the broad-spectrum aspartic acid protease inhibitor, pepstatin. This inhibitor was chosen as aspartic acid proteases are generally most active at acidic pH and therefore these enzymes were likely to dominate in the pH 3.5 assay conditions. In addition, our previous work with aspartic acid proteases in bovine secretory vesicles revealed that aspartic acid proteases had endoprotease activity and a preference for cleaving the peptide library between Phe-Phe residues (28), as was also found in **Figure 4A**.

In the presence of pepstatin, the intensity of all cleavage products was reduced and were often not detected above background (**Figure 5D**). These studies reveal that aspartic acid proteases are the dominant enzyme family in acidified wound fluid.

Using GKPILFFRL, the specific activity at pH 3.5 was quantified in wound fluid collected on Day 1, 3, 5, 7 and 10 and found to range between 0.1 and 0.16 fluorescence units/s/μg. In all samples this protease activity was reduced by 90% or more in the presence of pepstatin (**Figure 6A**) thereby confirming that the fluorogenic substrate is reporting on an aspartic acid protease. In addition, using this substrate, we showed that the specific activity was higher for the Day 10 sample relative to Day 5 wound fluid samples, as was seen with the MSP-MS assay.

**Figure 6.**
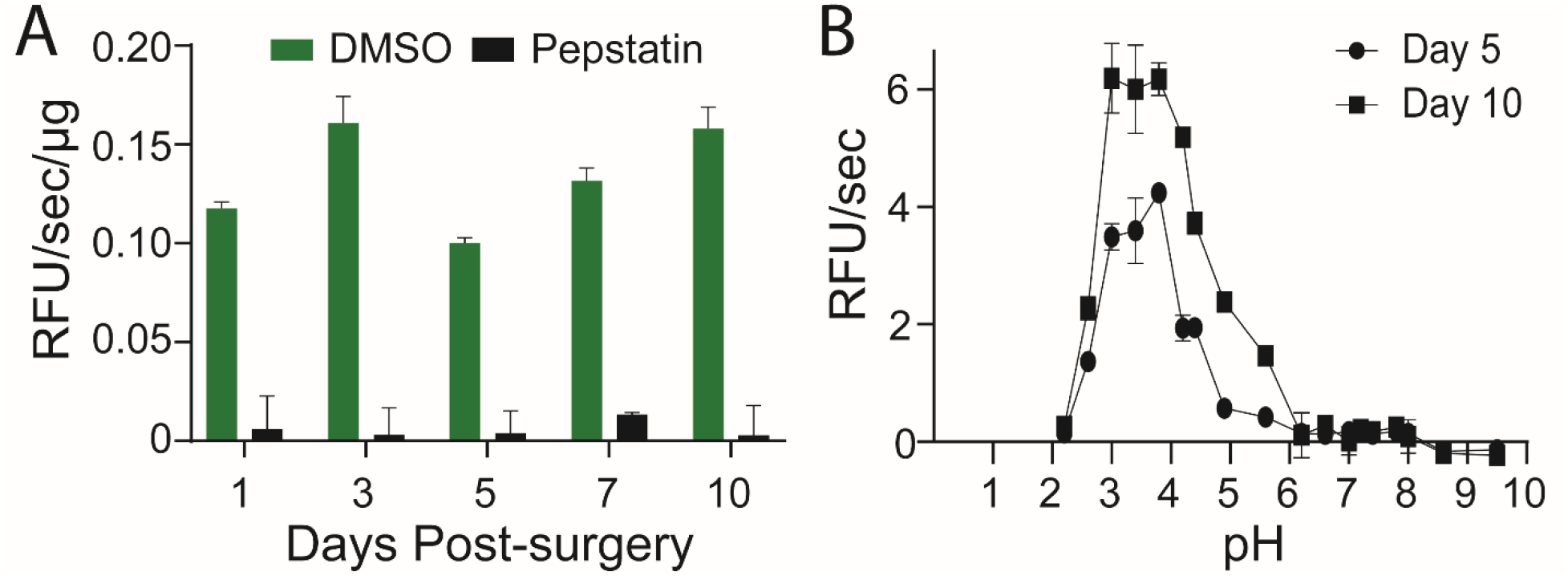
Mouse wound fluid protease activity **A.** Protease activity in acidified wound fluid (pH 3.5) from day 1 to 10 in the absence and presence of pepstatin. **B**. Superposition of pH curve of mice wound fluid from Day 5 and 10 post injury using GKPILFFRL substrate. Each datapoint corresponds to technical triplicates from one pooled wound fluid sample.

We next assayed Day 5 and Day 10 wound fluid across a wide pH range to determine the optimal environment that the aspartic acid proteases retain catalytic activity. Activity was low at pH 2.2 but increased steadily as the pH increased to 3.8. The maximum activity was found to be between pH 3.0 and pH 3.8. Above pH 3.8 activity decreased and no fluorescence change above background was detected at pH 6.2 and above (**Figure 6B**). This data reveals that the aspartic acid proteases present in wound fluid are optimally active between pH 3.0 and 3.8 of the wound fluid but are inactive at neutral pH. Therefore, these enzymes cannot contribute to wound proteolysis at neutral pH but are likely to be the dominant activity in acidified wounds.

To determine if the acid protease activity detected in murine wound fluid is equivalent in human samples, we obtained wound fluid from four patients of which one was an AIS patient (#55) and three were NM patients (#51, #57, and #59). Encouragingly, the reporter substrate for acid proteases in the murine samples (GKPILFFRL) was also cleaved by proteases in the patient samples (**Figure 7A**). Activity was highest at pH 3.5 and decreased as the pH was increased to pH 5.0 and pH 7.4. Using a panel of class-specific protease inhibitors, we showed that activity at pH 3.5 was inhibited only by the aspartic acid protease inhibitor, pepstatin and not by EDTA (metalloprotease inhibitor), Marimastat (MMP inhibitor), AEBSF (serine protease inhibitor), E64 (cysteine protease inhibitor) or carfilzomib (proteasome inhibitor). This data suggests that human wound fluid contains one or more aspartic acid proteases that are optimally active in acidic pH conditions and therefore are likely to function during CA treatment (**Figure 7A**). As a control, we also tested a human MMP-2 substrate (PLGLdAR) (29) in the same pH conditions and was found to be cleaved exclusively at pH 7.4 for patient samples 51, 55, and 59 (**Figure 7C**). For patient 57, cleavage of this substrate was also detectable at pH 5.5 and pH 3.5, albeit at a lower level than at pH 7.4. Using the same set of inhibitors, we show that cleavage of PLGLdAR is not inhibited by pepstatin but is inhibited by EDTA and Marimastat, confirming that activity is due to MMP-2 (**Figure 7D**).

**Figure 7.**
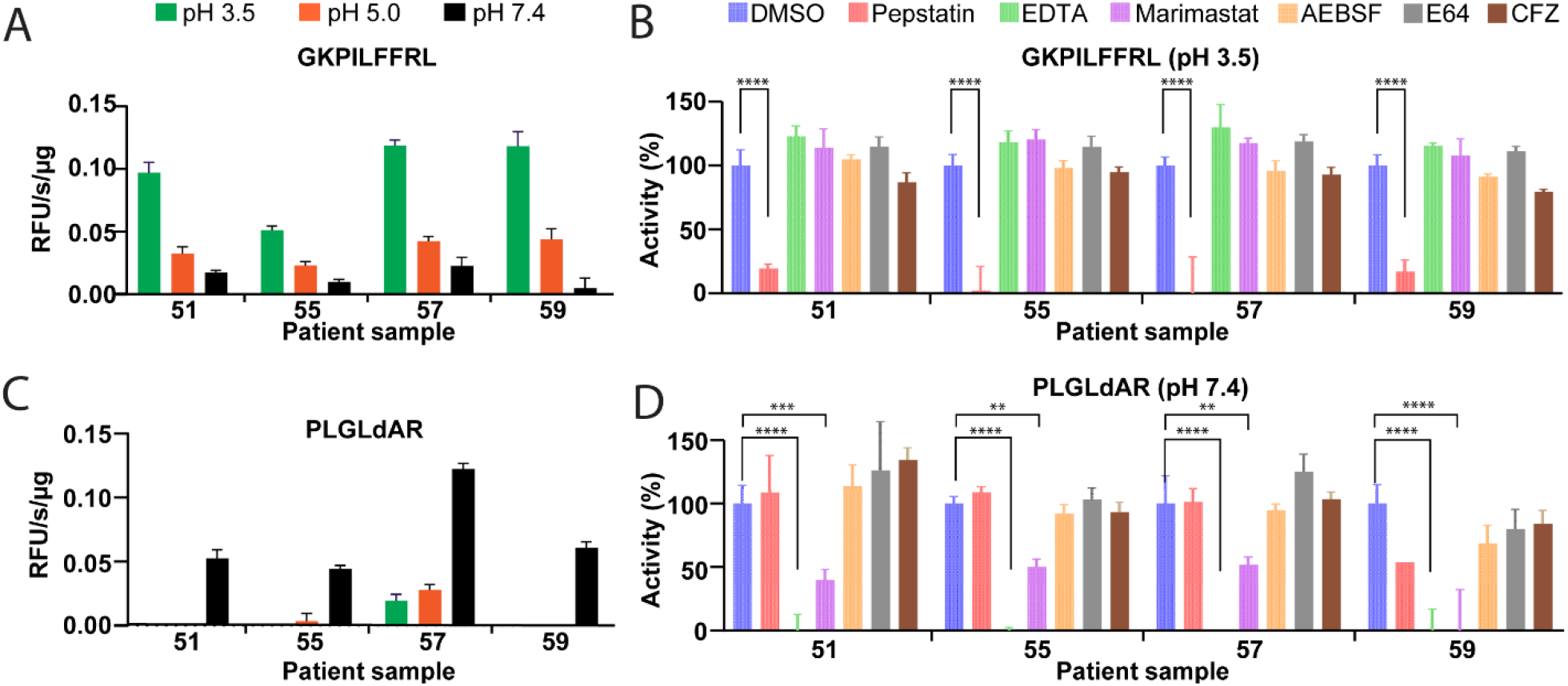
Human wound fluid protease activity **A**. Activity profile of wound fluid proteases using the GKPILFFRL substrate when assayed across different pH conditions. **B**. Inhibition profile of acid protease. Assays were performed in the absence and presence of the 10 µM pepstatin, 1 mM EDTA, 100 µM marimastat, 1 mM AEBSF, 10 µM E-64, and 10 µM CFZ inhibitors C. Activity profile of wound fluid proteases using the PLGLdAR substrate when assayed across different pH conditions. D. Inhibition profile of MMP-2 protease in wound fluid. Each data point corresponds to one assay performed in triplicate wells.

The two fluorogenic substrates GKPILFFRL and PLGLdAR are each selective for proteases that are active at either acid or neutral pH, respectively. We sought to find a reporter substate that is cleaved at both pH conditions so that we could directly compare the rate of cleavage. We screened a small library of in-house fluorescent peptides and determined that RPKPVEvWR was a substrate that could be cleaved by human wound fluid proteases at pH 3.5 and pH 7.4 (**Figure 8A**). This substrate was originally developed for cleavage by MMP-3 (30). For patient 51, 57 and 59 no cleavage of this substrate occurs at pH 5.0 indicating that the enzyme(s) responsible for cleaving this substrate at pH 7.4 are not active at pH 5.0. When assayed at pH 3.5, this substrate was cleaved by all wound fluid samples and this activity was inhibited by pepstatin, revealing that it was cleaved by the same enzyme that cleaves GKPILFFRL (**Figure 8B**). When assayed at pH 7.4, AEBSF significantly reduced activity in 3 out of 4 patients revealing that a serine protease was responsible for this enzyme activity. No reduction was seen in the presence of the MMP inhibitors, EDTA and Marimastat (**Figure 8C**) making it likely that MMP-3 is not active in these samples. Overall, this substrate is hydrolyzed equally well by serine proteases at neutral pH and aspartic acid proteases at pH 3.5.

**Figure 8:**
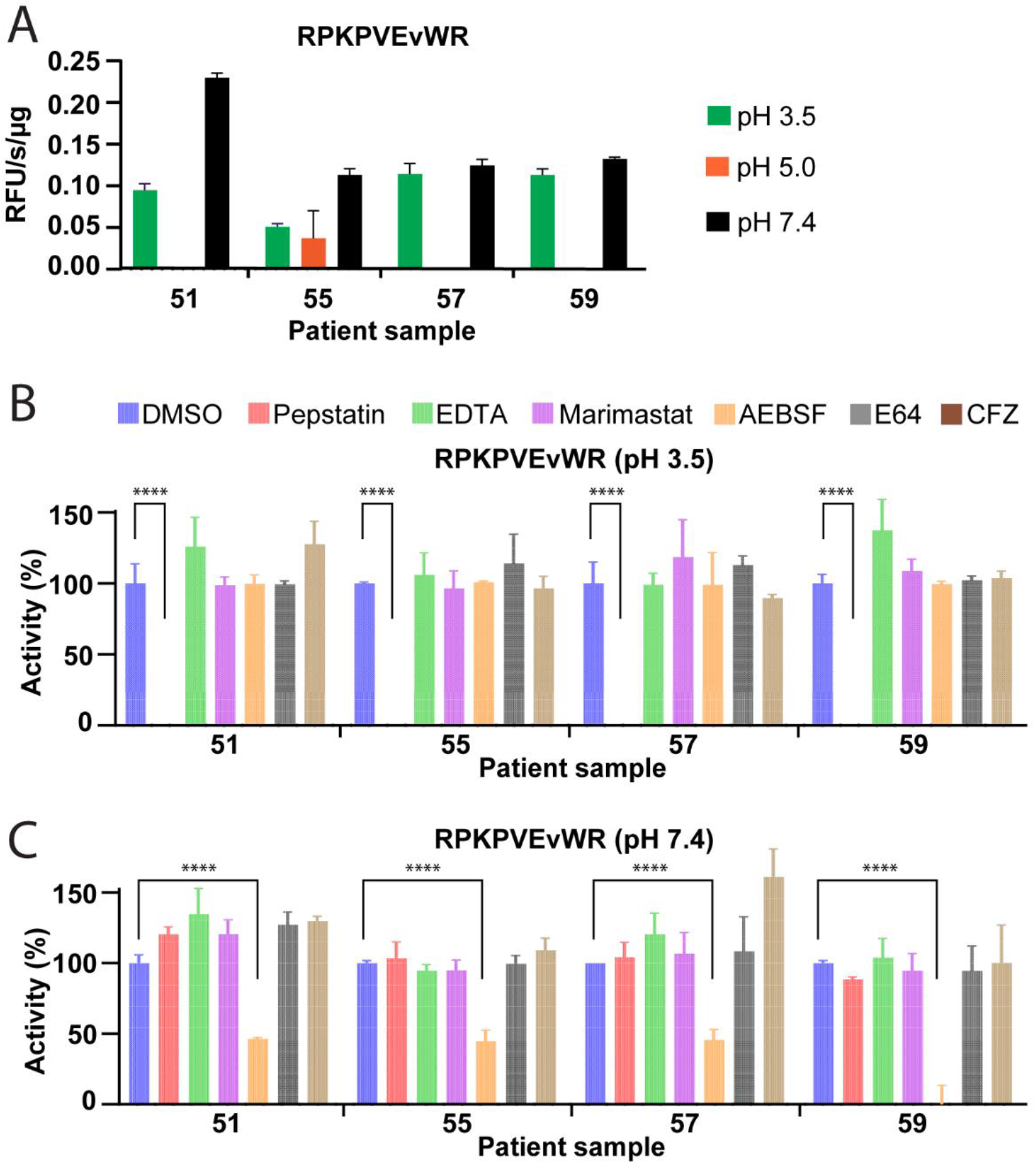
Protease activity assays of human wound fluid proteases using RPKPVEvWR as substrate. Assays were performed in the absence and presence of the 10 uM pepstatin, 1 mM EDTA, 100 µM marimastat, 1 mM AEBSF, 10 µM E-64, and 10 µM CFZ inhibitors at **A**. pH 3.5 and **B**. pH 7.4

The studies show that pH plays a key role in determining the active proteases in wound fluid and suggests that we can potentially improve wound healing rate and quality by carefully adjusting the pH of the wound using CA, acetic acid or other acidic reagents.

## Discussion

Our study revealed the presence of an aspartic acid protease in wound fluid that is active only under acidic pH conditions. This finding is important as it sheds light on the proteolysis that occurs when wounds are treated with CA and acetic acid. At neutral pH, achieving a balance between synthesis and breakdown of matrix components is crucial for wound healing. This balance requires the precise regulation of key proteases such as cathepsin G, elastase, plasmin, and MMP2 and proteases inhibitors such as serpins and TIMPs (1,5). Greener et al. demonstrated that these enzymes are optimally active from pH 7 to 9 which aligns with the pH range typically found in wounds. Acidification of wounds to pH 4 results in an 80% reduction in the activity of these enzymes (19) but our study reveals that a new set of proteases are likely to become active in these conditions.

The pH of the skin surface is typically between pH 4 and 6 and this acidity is crucial for maintaining a healthy stratum corneum and for epidermal barrier homeostasis (31). Owing to these conditions, it is not surprising that acid-acting proteases are present in the skin. Cathepsins L, B and D are all acid-acting proteases that are present in the epidermis and western blotting studies revealed that they were mostly present as inactive pro-enzymes (32). Proteomic studies also revealed that these enzymes are present in wound fluid (33). Therefore, acidification of wound fluid would likely result in activation of these enzymes.

When wound fluid was acidified to pH 3.5, we detected robust cleavage of a diverse peptide library and a resulting substrate preference for cleavage between hydrophobic amino acids. We have previously observed a similar profile when characterizing human cathepsin D using the same peptide library (34). Addition of pepstatin, a pan-aspartic acid inhibitor eliminated this activity revealing that that the enzyme responsible for this activity was likely to be cathepsin D. This observation was further supported by the efficient cleavage of a cathepsin D fluorogenic substrate in wound fluid following acidification to pH 3.5. Taken together, these data strongly indicate that cathepsin D in wound fluid is activated upon exposure to acid pH.

The protease activity that was detected in wound fluid at pH 5.0 may be due in part to cathepsin D, as this retains some activity up to pH 5.5. However, additional proteases are active at pH 5.0 which could be cathepsin B and L as these enzymes are optimally active from pH 4.5 to pH 6.0. Both enzymes are inactivated by the cysteine protease inhibitor, E64, and therefore future studies will be performed to determine if this inhibitor can uncover the role of cysteine proteases in mildly acidic conditions.

Studies have highlighted the advantageous role of an acidic environment in various stages of wound healing. CA treatment, for instance, is believed to facilitate wound healing by stimulating epithelialization through fibroblast proliferation, boosting local oxygen levels, and inhibiting microbial growth and virulence (10). Prolonged acidification of wound surfaces has also been linked to enhanced healing, as it augments cellular oxygen availability within the wound area (35). Furthermore, pH reduction in the wound environment correlates with heightened DNA synthesis (36), indicating a potential mechanism for accelerated tissue repair.

Studies have reported that cathepsin D activity increases at varying intervals post-skin injury. Notably, the topical application of pepstatin A effectively curtailed the rise in protein expression and cathepsin D activity, consequently delaying the repair of permeability barriers following disruption (37). These results suggest that, alongside neutral proteases, this aspartyl protease significantly contributes to the epidermal repair process post-injury.

In this study, we have revealed that cathepsin D is likely to be the dominant protease in acid-treated wounds. However, the current understanding of acid-mediated wound healing remains incomplete. More studies are needed to determine the optimal duration of treatment and the concentration and type of acid that is best to use to reach the target pH. Furthermore, the impact of acid treatment on surrounding healthy tissues requires thorough investigation. A deeper understanding of the biochemical mechanisms involved, and a rigorous evaluation of safety are essential before widespread clinical use of acid in wound healing.

## Acknowledgements

A.M.J would like to acknowledge the Carney Institute Innovation Awards, DARPA Director’s Award, NHLBI 3R01HL126887-02S1, NHLBI R01HL12688 and R01HL165259-01A1.

## Notes

### Competing Interest Statement

The authors have declared no competing interest.

